# DL4MicEverywhere: Deep learning for microscopy made flexible, shareable, and reproducible

**DOI:** 10.1101/2023.11.19.567606

**Authors:** Iván Hidalgo-Cenalmor, Joanna W Pylvänäinen, Mariana G Ferreira, Craig T Russell, Ignacio Arganda-Carreras, AI4Life Consortium, Guillaume Jacquemet, Ricardo Henriques, Estibaliz Gómez-de-Mariscal

## Abstract

Deep learning has revolutionised the analysis of extensive microscopy datasets, yet challenges persist in the widespread adoption of these techniques. Many lack access to training data, computing resources, and expertise to develop complex models. We introduce DL4MicEverywhere, advancing our previous ZeroCostDL4Mic platform, to make deep learning more accessible. DL4MicEverywhere uniquely allows flexible training and deployment across diverse computational environments by encapsulating methods in interactive Jupyter notebooks within Docker containers –a standalone virtualisation of required packages and code to reproduce a computational environment–. This enhances reproducibility and convenience. The platform includes twice as many techniques as originally provided by ZeroCostDL4Mic and enables community contributions via automated build pipelines. DL4MicEverywhere empowers participatory innovation and aims to democratise deep learning for bioimage analysis.

## Introduction

Deep learning enables the transformative analysis of large multidimensional microscopy datasets, but barriers remain in implementing these advanced techniques (3, 4). Many researchers lack access to sufficient annotated data, highperformance computing resources, and expertise to develop, train, and deploy complex deep-learning models. In recent years, several approaches have been developed to democratise the usage of deep learning for microscopy (4). Multiple tools, such as BioImage.io, facilitate sharing and reusing broadly useful, previously trained deep learning models, distributing them as one-click image analysis solutions (1, 5). Yet often, deep learning models need to be trained or finetuned on the end user dataset to perform well (1, 4, 6). We previously released ZeroCostDL4Mic (2), an online platform relying on Google Colab that helped democratise deep learning by providing a zero-code interface to train and evaluate models capable of performing various bioimage analysis tasks, such as segmentation, object detection, denoising, super-resolution microscopy, and image-to-image translation. Here, we introduce DL4MicEverywhere, a major advancement of the ZeroCostDL4Mic (2) framework (**Fig.1**). DL4MicEverywhere allows users the flexibility to train and deploy their models across various computational environments, including Google Colab, their own computational resources (*e*.*g*., desktop or laptop), or high-performance computing systems. This flexibility is made possible by enclosing each deep learning technique in an interactive Jupyter notebook, which is then contained in a Docker (7)-based environment. This enables users to install and interact with deep learning techniques easily. Incorporating crossplatform containerisation technology boosters the long-term platform’s stability and reproducibility and enhances user convenience (8).

**Fig. 1.**
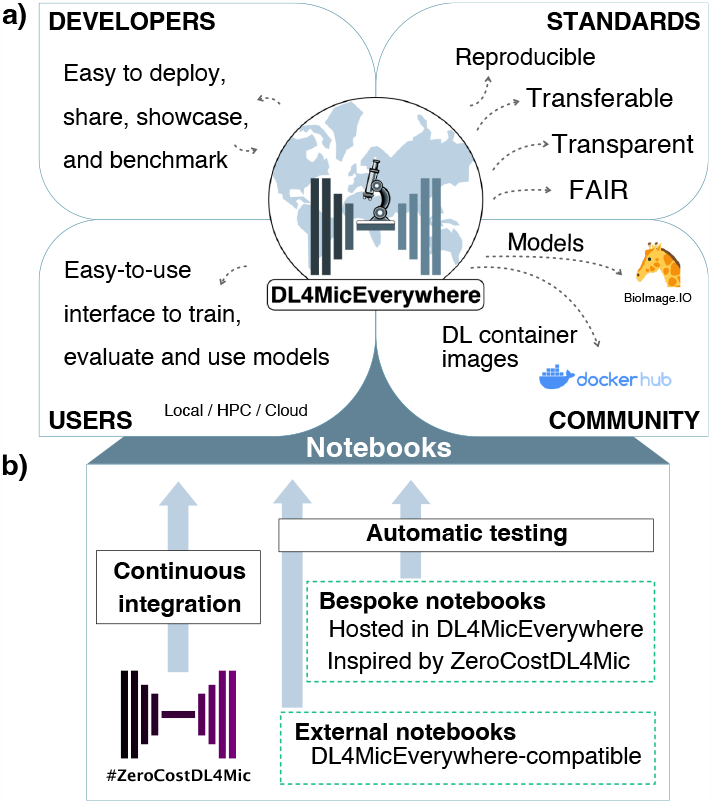
DL4MicEverywhere platform. a) DL4MicEverywhere eases deep learning workflow sharing, deployment, and showcasing by providing a user-friendly interactive environment to train and use models. Enabling cross-platform compatibility ensures deep-learning model training reproducibility. DL4MicEverywhere contributes to deep learning standardisation in bioimage analysis by promoting transferable, FAIR, and transparent pipelines. The platform exports models compatible with the BioImage Model Zoo(1) and populates the Docker hub with free and open source (FOSS) container images that developers can reuse, incrementing the list of available workflows. b) DL4MicEverywhere accepts three types of notebook contributions: ZeroCostDL4Mic(2) notebooks, bespoke notebooks inspired by ZeroCostDL4Mic(2), and notebooks hosted in external repositories that are compliant with our format. These contributions are automatically tested to ensure the correct requirements and format.

**Fig. 2.**
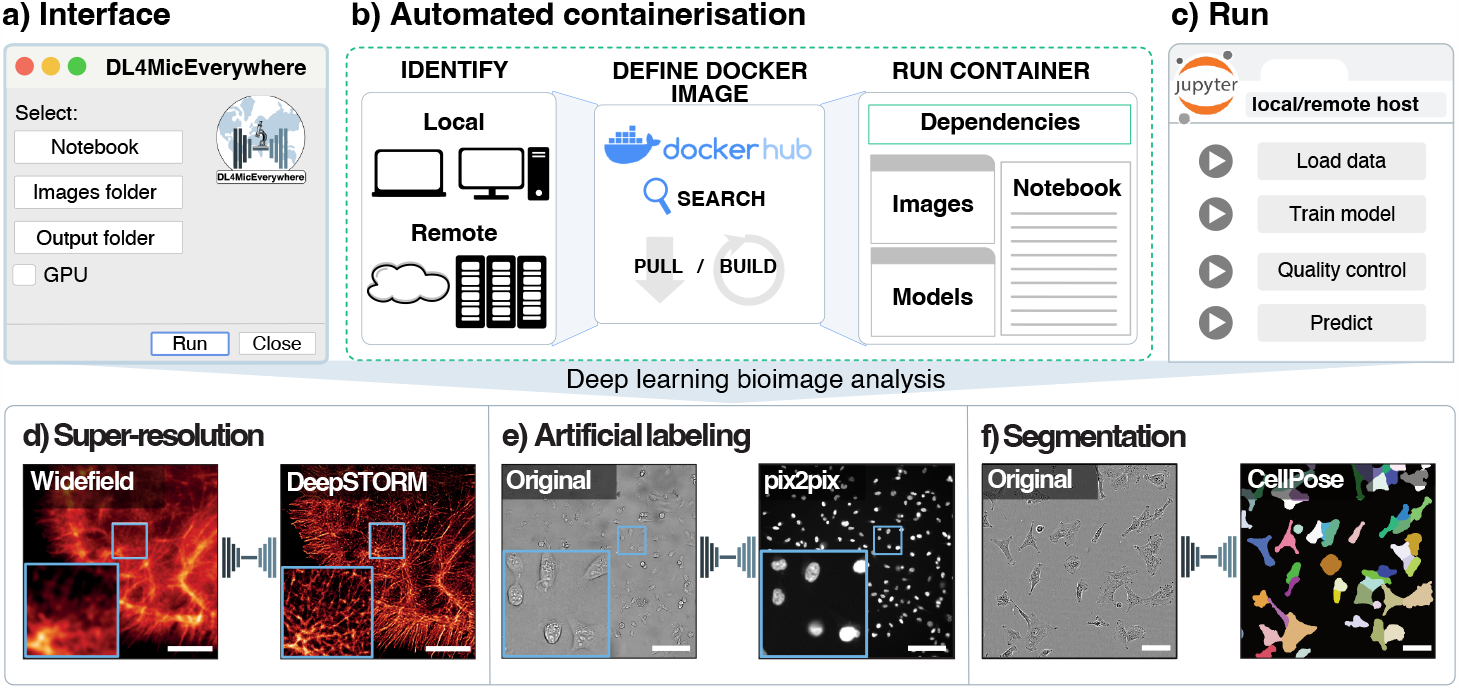
a) When running DL4MicEverywhere, the user interacts with an interface to choose a notebook, image and output folder, and choose a GPU running model if possible. b) DL4MicEverywhere will automatically identify the system architecture and requirements to build a Docker container image. If the image is not available in the Docker hub, it is built in the user’s machine. This image is used to create a Docker container: a functional instance of the image that gathers the code environment to use the chosen notebook. c) A Jupyter lab session is launched inside the Docker container to train, evaluate or use the chosen deep learning model. DL4MicEverywhere notebooks are also interactive and equivalent to ZeroCostDL4Mic (2) notebooks. d-f) DL4MicEverywhere enables the use of the same notebooks in different local or remote infrastructures such as workstations, the cloud or high-performance computing clusters. This is, researchers could run exactly the same d) super-resolution, e) artificial-labelling or g) segmentation pipelines, among many others, in different systems.

## Results

DL4MicEverywhere introduces a novel and user-friendly graphical interface that enables users to easily access and launch a comprehensive collection of interactive Jupyter notebooks. Each notebook comes packaged into a Docker container with all necessary software dependencies, as illustrated in **Fig. 2a-c**.

DL4MicEverywhere has gone beyond simply containerising notebooks, providing a zero-code interface that handles all behind-the-scenes complexities. Users are not required to deal with the intricacies of Docker or configuring deep learning frameworks. The intuitive interface abstracts away these technical details, while the Docker encapsulation provides a standardised and rich environment for executing advanced techniques reliably (**Figure 2b**). Researchers can select a notebook, choose computing resources, and run the corresponding deep learning-powered analysis with just a few clicks. The platform handles deploying the encapsulated coding environment seamlessly in the background. This allows users to train and apply models on various computing resources they control, eliminating reliance on third-party platforms. Furthermore, researchers can launch a notebook on local or remote systems with GPU acceleration on clusters whenever available, without worrying about complex software dependencies, docker container management or losing access to deep-learning frameworks (**Fig. 2d-f**). DL4MicEverywhere offers twice the number of deep learning approaches than what was initially available in ZeroCostDL4Mic. The platform is designed to encourage sharing and reuse of models via the BioImage Model Zoo. DL4MicEverywhere’s infrastructure is strengthened by automated build pipelines (9), which allows for the seamless integration of new trainable models contributed by the community (10–13) (as shown in **Fig. 1b**). These contributions are further facilitated through user-friendly templates, allowing new notebooks to be added independently of the original ZeroCostDL4Mic framework. By empowering participatory innovation in an open and flexible platform, DL4MicEverywhere aims to make deep learning more accessible for bioimage analysis. Developers can share a notebook based on our template and metadata for their method, and DL4MicEverywhere handles the testing and building of fully documented and open-source containerisation. Note that notebook containerisation allows others to reliably replicate analyses and build on the latest methods. The highly flexible nature of Docker containers encapsulating notebooks enhances long-term reproducibility across operating systems and computing environments. Researchers can easily share not just code, but the full software environment required to run it reliably. This reusable encapsulation empowers others to replicate analysis, evaluate methods, and build on research.

## Discussion

Deep learning is revolutionising microscopy through datadriven analysis and discovery (14). However, significant barriers persist in accessing these advanced techniques, including a lack of training data, computing resources, and expertise (4, 6, 14). Proprietary platforms create technological and cultural obstacles, while complex workflows impede adoption by non-experts. DL4MicEverywhere is an initiative that aims to make deep learning accessible to everyone by providing a flexible and community-driven platform. Encapsulating software in Docker containers makes it possible to integrate new methods and enrich the microscopy community through participatory innovation. Intuitive graphical user interfaces also lower the barriers to entry, making it easier for non-experts to use the platform. Users can rely on shared techniques while customising models across diverse hardware, retaining control over data and analysis. The platform will particularly be useful with the increasing development and use of cutting-edge foundation models (15). By bundling these sophisticated models into shareable containers, researchers can customise and exploit them in their microscopy applications. DL4MicEverywhere also simplifies complex deep learning workflows for nonexperts through automated pipelines, and is optimised for use with local computational resources, high-performance computing, and cloud-based solutions. This flexibility is precious for 1) sensitive biomedical data, where privacy risks may limit reliance on public cloud platforms, and 2) continuously scaling data such as time-lapse volumetric images or high-throughput high-content imaging data, where storage, dissemination and access rely on institutional infrastructures with specific data sharing protocols. DL4MicEverywhere also adheres to FAIR principles, enhancing discoverability and interoperability. We expect DL4MicEverywhere to represent an important step towards reliable, transparent, and participatory artificial intelligence in microscopy.

## Code availability

The source code, documentation, and tutorials for DL4MicEverywhere can be found at https://github.com/HenriquesLab/DL4MicEverywhere. DL4MicEverywhere is made available under the Creative Commons CC-BY-4.0 license.

## ACKNOWLEDGEMENTS

I.H.C., M.G.F., C.T.R., R.H., and E.G.M. received funding from the European Commission through the Horizon Europe program (AI4LIFE project with grant agreement 101057970-AI4LIFE, and RT-SuperES project with grant agreement 101099654-RT-SuperES to R.H.). I.H.C., M.G.F., E.G.M. and R.H. also acknowledge the support of the Gulbenkian Foundation (Fundação Calouste Gulbenkian) and the European Research Council (ERC) under the European Union’s Horizon 2020 research and innovation programme (grant agreement No. 101001332 to R.H.). Funded by the European Union. Views and opinions expressed are however those of the authors only and do not necessarily reflect those of the European Union. Neither the European Union nor the granting authority can be held responsible for them. This work was also supported by the European Molecular Biology Organization (EMBO) Installation Grant (EMBO-2020-IG-4734 to R.H.), the EMBO Postdoctoral Fellowship (EMBO ALTF 174-2022 to E.G.M.), the Chan Zuckerberg Initiative Visual Proteomics Grant (vpi-0000000044 with DOI:10.37921/743590vtudfp to R.H.) and the Chan Zuckerberg Initiative DAF, an advised fund of Silicon Valley Community Foundations (Chan Zuckerberg Initiative Napari Plugin Foundations Grant Cycle 2, NP2-0000000085 granted to R.H.). R.H. also acknowledges the support of LS4FUTURE Associated Laboratory (LA/P/0087/2020). This work is partially supported by grant GIU19/027 (to I.A.C.) funded by the University of the Basque Country (UPV/EHU), grant PID2021-126701OB-I00 (to I.A.C.) funded by the Ministerio de Ciencia, Innovación y Universidades, AEI, MCIN/AEI/10.13039/501100011033, and by “ERDF A way of making Europe” (to I.A.C.). This study was also supported by the Academy of Finland (338537 to G.J.), the Sigrid Juselius Foundation (to G.J.), the Cancer Society of Finland (Syöpäjärjestöt; to G.J.), and the Solutions for Health strategic funding to Åbo Akademi University (to G.J.). This research was supported by InFLAMES Flagship Programme of the Academy of Finland (decision number: 337531). We would like to thank Amin Rezaei, Ainhoa Serrano, Pablo Alonso, Urtzi Beorlegui, Andoni Rodriguez, Erlantz Calvo, Soham Mandal, and Virginie Uhlmann for their contributions to the ZeroCostDL4Mic notebook collection.

## Authors of the AI4Life Horizon Europe program consortium

**Arrate Muñoz-Barrutia**

Bioengineering Department, Universidad Carlos III de Madrid, Leganes, Spain; Instituto de Investigación Sanitaria Gregorio Marañón, Madrid, Spain

**Beatriz Serrano-Solano**

Euro-BioImaging ERIC Bio-Hub, European Molecular Biology Laboratory (EMBL) Heidelberg, Heidelberg, Germany

**Caterina Fuster Barcelo**

Bioengineering Department, Universidad Carlos III de Madrid, Leganes, Spain; Instituto de Investigación Sanitaria Gregorio Marañón, Madrid, Spain

**Constantin Pape**

Georg-August-University Göttingen, Institute of Computer Science, Göttingen, Germany

**Emma Lundberg**

Department of Pathology, Stanford University School of Medicine, Stanford, CA, USA; Science for Life Laboratory, School of Engineering Sciences in Chemistry, Biotechnology and Health, KTH Royal Institute of Technology, Stockholm, Sweden; Department of Bioengineering, Stanford University, Stanford, CA, USA

**Florian Jug**

Fondazione Human Technopole, Milan, Italy

**Joran Deschamps**

Fondazione Human Technopole, Milan, Italy

**Matthew Hartley**

European Molecular Biology Laboratory, European Bioinformatics Institute, EMBL-EBI, Wellcome Genome Campus, Cambridge, UK

**Mehdi Seifi**

Fondazione Human Technopole, Milan, Italy

**Teresa Zulueta-Coarasa**

European Molecular Biology Laboratory, European Bioinformatics Institute, EMBL-EBI, Wellcome Genome Campus, Cambridge, UK

**Vera Galinova**

Fondazione Human Technopole, Milan, Italy

**Wei Ouyang**

Department of Applied Physics, Science for Life Laboratory, KTH Royal Institute of Technology, Stockholm, Sweden

## Methods

### DL4MicEverywhere Platform Implementation

The core DL4MicEverywhere platform was implemented in Bash packaging and managing Python notebook workflows through Docker containers. An overview of the key technical components is provided below.

### Docker Containerization

Each notebook is encapsulated into a Docker container, including all dependencies required for smooth runtime (Docker v24.0.5, Docker Inc.). These containers are functional instances of Docker images –software units that contain the virtualisation of a specific computational environment, with all the specified dependencies and packages included. Images were built from Ubuntu (v20.04/22.04) base images, with optional Nvidia CUDA support for GPU acceleration. Python (v3.7/3.8/3.10), deep learning packages (TensorFlow, Keras or Pytorch), and notebook packages were installed according to the requirements into the containers. Unique containers were constructed for each notebook using a parameterised Docker file build process, taking metadata like notebook URL and software versions as input. These images are uploaded to Docker hub so they can be distributed as free and open source (FOSS) and belong to the Open Container Initiative (OCI) (https://opencontainers.org/).

### Launch Script and GUI

A Bash shell script launch.sh was implemented to manage the building, running, and monitoring of the notebook containers based on user input. Key functions included argument parsing, installation checking, Docker image building, and Jupyter Lab invocation within the container. A graphical user interface was additionally created using Wish (a Tcl/Tk application) to enable intuitive notebook and parameter selection through a desktop window. This is invoked by the launch script and passed user selections.

### Configuration Metadata

Inspired by the BioImage Model Zoo (1) specifications, notebook container construction was driven by human-readable YAML configuration files specifying necessary build metadata for each notebook, including the URL of the notebook itself, Python requirements, and Docker parameters. These configurations were loaded by the launch script when initialising a container. This format establishes a basis for a seamless connection with the BioEngine of the Zoo.

### Testing and Deployment Pipelines

GitHub Actions workflows were implemented to automatically build and publish container images for each new notebook, handling testing across platforms like AMD64 and ARM64. Images were versioned based on notebook metadata and published to DockerHub for distribution. Strict conventions enforced by templates facilitated notebook contributions from the community. These contributions are further checked via GitHub Actions to assert that they follow the specified format with valid URLs and that it is possible to build a Docker image.

### Jupyter Notebooks and Widgets

Notebooks were adapted from the ZeroCostDL4Mic Colab format to interactive Jupyter notebooks leveraging *ipywidgets* for a simplified user interface requiring no coding. Parameters could be configured via graphical elements rather than edits to code.

